# OCTA observation of changes in macular vessel density in diabetic patients and its correlation with diabetic retinopathy staging : A cross-sectional study

**DOI:** 10.1101/2022.02.07.479361

**Authors:** Deng Yu, Jie Chuanhong, Wang Jianwei, Liu Ziqiang, Li Yuanyuan

## Abstract

**Objective:** To investigate the association between disease progression and OCTA vessel density and other indices in patients with diabetic retinopathy.

**Methods:** Participants were selected with the following criteria: 63 patients (100 eyes) diagnosed with type 2 diabetes mellitus, which included 44 patients (72 eyes) with diabetic retinopathy and 19 patients (28 eyes) with type 2 diabetes mellitus and non-diabetic retinopathy (NDR), who were seen at the Eye Hospital China Academy of Chinese Medical Sciences from September 2020 to July 2021. All patients underwent OCTA examination, and FAZ, PERIM, AI, FD, SVD, DVD and other indices were counted.

**Results:** (1) The correlation coefficients of SVD, paracentric SVD, DVD, paracentric DVD and DR processes were: -0.525, -0.586, -0.323, and -0.424 (*P*< 0.05), respectively, and all were moderately negatively correlated. (2) The correlation coefficients of FAZ and PERIM with DR process were: -0.031, 0.084 (*P*>0.05), respectively, and not correlated. The correlation coefficients of AI and FD with DR process were: 0.307, −0.459 (*P*<0.05), and with moderate positive and negative correlations, respectively. (3) The correlation coefficients of FAZ, PERIM, AI and FD with age were: -0.124, -0.052, 0.113, -0.170 (*P*>0.05), and no correlation, respectively.

**Conclusion:** The disease progression of DR was moderately correlated with OCTA superficial vessel density and deep vessel density; and moderately correlated with AI and FD. OCTA could assist in the assessment of DR disease progression.

## 1. Introduction

Diabetic retinopathy (DR) is one of the most common microvascular complications of diabetes mellitus and is one of the leading causes of vision loss in patients with diabetes mellitus (DM). Approximately 35% of diabetic patients are affected by DR^[1]^. The pathological process of DR includes pericyte loss, endothelial cell proliferation, basement membrane thickening, canalicular stenosis and occlusion, retinal ischemia and hypoxia, neovascularization, traction retinal detachment, etc.^[2]^. The detection and quantification of early retinal microvascular damage in DM patients could provide a better understanding of the pathogenesis and predict the progression of DR.

Optical coherence tomography angiography (OCTA) is a non-invasive vascular imaging technique, which allows the rapid acquisition of different levels of vessel angiography images of the retinal and choroidal vessels without the use of contrast agents^[3]^. OCTA can be used to identify non-perfused areas, microaneurysms, and neovascularization, vessel density (VD), etc. These parameters allow quantification and statistical analysis of the macular peripheral microvascular damage and the area of the foveal avascular zone (FAZ)^[4]^. OCTA has better visualization^[5]^ and can detect microscopic neovascularization that is difficult to detect on clinical examination, assess the extent^[6]^and activity^[7]^ of neovascularization in patients with proliferative diabetic retinopathy (PDR), and evaluate the changes in retinal neovascularization after anti– vascular endothelial growth factor (anti-VEGF) and retinal photocoagulation treatments^[8]^. Although there have been numerous studies on OCTA in recent years, fewer studies have addressed its correlation with DR progression, and there were differences between studies addressing the same indices. In this study, conducted in Chinese patient, we collected the OCTA findings from patients with different stages of DR, analyzed their FAZ area, vessel density in the different areas of superficial and deep capillaries, and evaluated the correlation between them and DR staging to elucidate the changes in vessel density in different levels and areas of OCTA during DR progression, and to provide assistance for the clinical application of OCTA for the observation and detection of DR disease and progression.

## 2. Materials and methods

### 2.1 Patient source

63 patients (100 eyes) diagnosed with type 2 diabetes mellitus, who attended the Eye Hospital China Academy of Chinese Medical Sciences from September 2020 to July 2021. There were 28 male patients (46 eyes) and 35 female patients (54 eyes) with a mean age 61.2 years.The ethical board of the China Academy of Chinese Medical Sciences and Eye Hospital China Academy of Chinese Medical Sciences approved this study. The work described has been carried out in accordance with the Code of Ethics of the Declaration of Helsinki.

### 2.2 Diagnostic criteria

We retrospectively identified patients with diabetic retinopathy according to the new 2002 International Clinical Diabetic Retinopathy (ICDR) severity scale: (1) No significant retinopathy: no abnormalities; (2) mild non-proliferative diabetic retinopathy (NPDR): microaneurysm only; (3) moderate NPDR: microaneurysm with milder than severe NPDR manifestations; (4) severe NPDR: any 1 of the following changes, but without NPDR manifestations: more than 20 intraretinal hemorrhages in any quadrant; venous beading in more than two quadrants; and significant intraretinal microvascular abnormalities in 1 quadrant; (5) Proliferative diabetic retinopathy (PDR): Presence of 1 or more of the following changes: neovascularization, vitreous hemorrhage, or preretinal hemorrhage.

### 2.3 Inclusion criteria for the observation group

(1) Those with a clear history of type 2 diabetes mellitus; (2) those who could cooperate in completing the OCTA examination.

### 2.4 Exclusion criteria

(1) Previous history of retinal laser photocoagulation; history of intraocular surgery, such as intravitreal injection, vitrectomy, etc. in the past 3 months; (2) history of age-related macular degeneration, high myopia, macular fissure, rhegmatogenous retinal detachment, vitreomacular traction syndrome, glaucoma, optic nerve atrophy, optic neuritis and other diseases affecting the function of the retina and optic nerves; (3) suffering from other systemic diseases that may affect ocular function; (4) cataract, vitreous hemorrhage and other reasons that render OCTA results unrecognizable.

### 2.5 Research methodology

#### 2.5.1 Confirmation of DR diagnosis and staging

The patient’s eye fundus was examined by 2 physicians, using a 90D ophthalmoscope after mydriasis, and determine the DR staging. When the results of the 2 physicians were inconsistent, the third physician re-examined the patient’s colored fundus photographic examination and determined the patient’s stage. Finally, the 3 physicians jointly decided the staging of the patient.

#### 2.5.2 OCTA examination

1 designated technologist performed OCTA examinations, using OCTA (Optovue. Fremont, CA, USA). Retina 3.0 mode (i.e., macular area 3 × 3 mm range scan mode) was used for arch ring observation: arch ring area, perimeter, acircularity index (AI), and vessel density within 300 μm outside of the arch ring. Central retinal thickness, superficial and deep retinal capillary vessel density. A 840 nm light source with a bandwidth of 50 nm was used, and a single image acquisition consisted of 1 horizontal scan superimposed on 1 vertical scan.

#### 2.5.3 Observational indices

VD in different regions such as FAZ, PEIM, AI, FD, SVD, and foveal and parafoveal DVD. Visual acuity was not included as an observational index in this study, because visual acuity is easily affected by lens opacity and other factors such as refractive interstitial and macular edema.

### 2.6 Statistical methods

SPSS 23.0 statistical software was used for data analysis. The results were expressed as M(_25_P^P_75_) for patient data, using the Kruskal-Wallis (K-W) test for normality, and non-parametric tests. And the non-parametric Mann-Whitney U test was used for comparison of data between groups. The correlation between different stages of DR and OCTA examination of FAZ, PERIM, AI, FD, and VD in different regions, and the correlation of age with FAZ, PERIM, AI, and FD were analyzed using Spearman’s test. The results were expressed using correlation coefficients.

### 2.7 Ethical consideration

This study was a retrospective study approved by the Eye Hospital China Academy of Chinese Medical Sciences. There were no ethical issues in conducting this study because it was a retrospective study targeting to find the relationship between OCTA and diabetic retinopathy, and the authors declared that all methods were performed in accordance with the relevant guidelines and regulations (Declaration of Helsinki).

## 3. Results

### 3.1 Basic information of patients

A total of 72 eyes of patients with diabetic retinopathy were included in this study based on the inclusion and exclusion criteria of this study, including mild or moderate NPDR: 31 eyes, aged 62.0 (58.0^68.0) years-old; severe or PDR: 41 eyes, aged 59.0 (52^64.0.0) years-old; and NDR patients: 28 eyes, aged 63.5 (62.0^69.8) years-old. There was no missing data. Differences were not statistically significant, using the Mann-Whitney U test.

### 3.2 OCTA results in patients with different staging

The K-W testing method was used to test whether the data conformed to a normal distribution (refer to “Statistics in Medicine (4th edition)” by Sun Zhenqiu et al.), and the results showed that, with the exception of FAZ, foveal DVD, PERIM, and central SVD conformed to a normal distribution, and a nonparametric test was used for the data. In each region of the SVD, the VD of patients in the severe or PDR group was lower than that of the mild or moderate group, the VD of patients in the mild or moderate group was lower than that of the NDR group in all regions, except the fovea, and in each region of the DVD, the VD of patients in the severe or PDR group was higher than that of the mild or moderate group, except for on the temporal side, and the VD of patients in the mild or moderate group was higher than that of the NDR group except for on the temporal side. The mean values were compared pairwise, and the differences between the groups were statistically significant (*P*<0.05), except for the SVD center and DVD center, suggesting that the indices were significantly different and that correlation analysis of multiple factors could be performed.

**Table 1:** Results of SVD rank sum test by region in patients with different stages.

| Superficial | SVD | Central | Paracentral | T | S | N | I |
| --- | --- | --- | --- | --- | --- | --- | --- |
| NDR | 44.8<br>(40.8^46.4) | 11.1<br>(6.2^18.9) | 47.0<br>(43.7^49.3) | 45.9<br>(41.8^49.1) | 48.6<br>(45.9^51.8) | 46.4<br>(43.9^48.0) | 49.2<br>(43.5^51.9) |
| Light/moderate | 42.3<br>(38.6^44.8)<br>* | 14.8<br>(9.7^22.4)<br>) | 45.0<br>(40.2^47.5)<br>* | 43.8<br>(40.0^46.5)<br>* | 44.6<br>(39.3^48.4)<br>* | 44.1<br>(37.9^48.0)<br>* | 47.0<br>(41.9^48.6)<br>* |
| Severe/PDR | 35.3<br>(.032^39.3)<br>*# | 11.4<br>(.08^16.2)<br>) | 37.0<br>(34.3^42.2)<br>*# | 36.2<br>(.033^41.6)<br>*# | 38.1<br>(35.3^43.2)<br>*# | 36.8<br>(33.6^41.9)<br>*# | 38.8<br>(33.9^41.0)<br>*# |
| Chi-square | 38.568 | 2.292 | 36.413 | 27.813 | 34.808 | 32.919 | 33.214 |
| P | 0.000 | 0.318 | 0.000 | 0.000 | 0.000 | 0.000 | 0.000 |

\*In comparison with NDR, $P < 0.05$ ; # in comparison with mild/moderate NPDR, $P < 0.05$ .

**Table 2:**
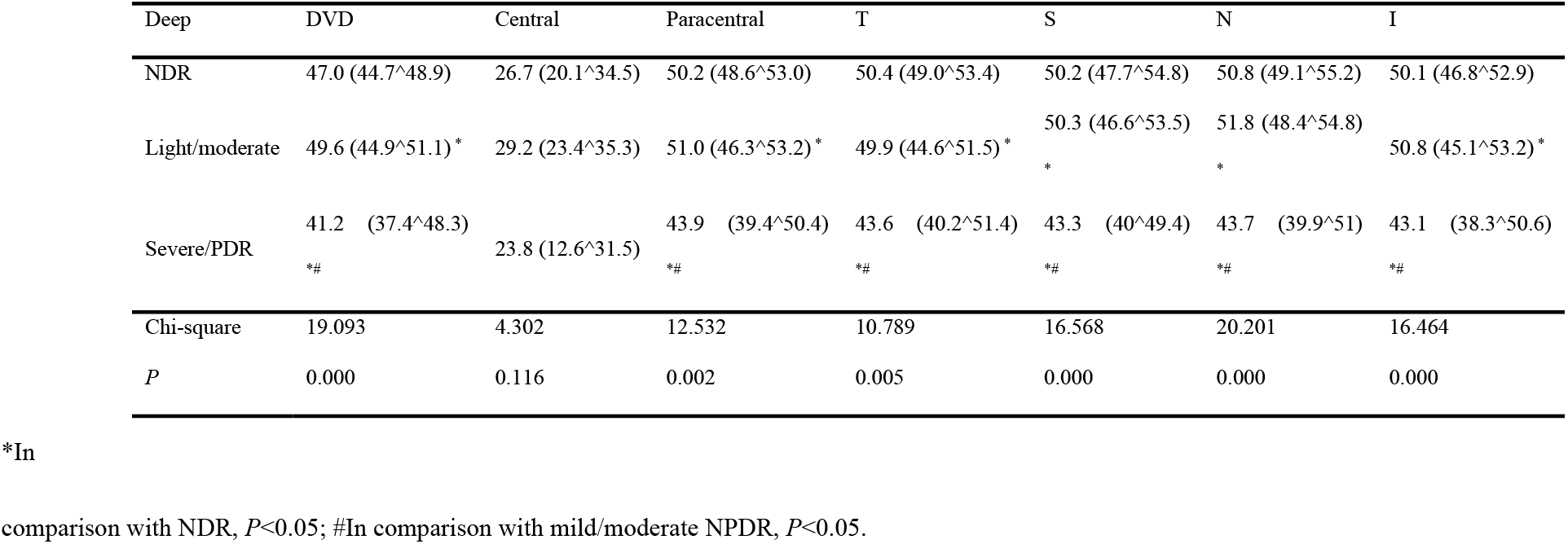
Results of DVD rank sum test by region in patients with different stages.

### 3.3

Correlation analysis was performed using Spearman’s test for overall SVD, paracentric SVD, temporal SVD, superior SVD, nasal SVD, and inferior SVD in patients with NDR, mild or moderate NPDR, moderate NPDR, and PDR, with correlation coefficients of −0.625, −0.661, −0.524, −0.587, −0.590, and −0.554, respectively (*P*<0.05), and correlation analyses for DVD, paracentral DVD, temporal DVD, superior DVD, nasal DVD, and temporal DVD derived correlation coefficients of: −0.340, −0.335, −0.292, −0.336, −0.407, and −0.344 (*P*<0.05), respectively. Correlation analysis was not performed, because the difference between central SVD, and central DVD was not statistically significant.

### 3.4 FAZ, PERIM, FD, AI and other findings in patients with different staging

**Table 3:** Results of FAZ, PERIM, FD and AI rank sum test.

|  | FAZ | PERIM | AI | FD |
| --- | --- | --- | --- | --- |
| NDR | 0.36 (0.3<sup>0.47</sup>) | 2.40 (2.13<sup>2.74</sup>) | 1.13 (1.1<sup>1.17</sup>) | 47.28 (44.18<sup>50.59</sup>) |
| Light/moderate | 0.34 (0.23<sup>0.42</sup>) | 2.47 (1.95<sup>2.78</sup>) | 1.17 (1.14<sup>1.22</sup>) | 46.97 (42.76<sup>49.82</sup>) |
| Severe/PDR | 0.31 (0.24<sup>0.58</sup>) | 2.43 (.002<sup>3.26</sup>) | 1.18 (1.14<sup>1.26</sup>) | 39.28 (33.72<sup>42.96</sup>) |
| $\chi^2$ | 0.820 | 0.606 | 8.935 | 28.050 |
| $P$ | 0.664 | 0.739 | 0.011 | 0.000 |

### 3.5 Correlation between DR progression with AI, PERIMFD and FAZ

Correlation analysis was performed using Spearman’s test for AI and FD in patients with NDR, mild or moderate NPDR, moderate NPDR, and PDR, and the correlation coefficients were: 0.280, −0.494, respectively. The data were all *P*<0.05, suggesting that the differences were statistically significant.

### 3.6 Differences in vessel imaging in patients with different stages of DR

Superficial OCTA flow imaging maps were observed in patients with different DR staging. In contrast to patients with NDR, patients with mild/moderate NPDR began to show small patches of nonperfused areas distributed along the pericentral sulcus, with roughly normal locations away from the central sulcus. Patients with severe NPDR became significantly more numerous than those with mild/severe NPDR or NDR without areas of concern, connected by small nonperfused areas in patches, whose vascular endings were seen to be tortuously extended and whose peripheral vessels began to be involved. Patients with PDR had significantly fewer microvessels, and non-perfusion zones in all quadrants were divided by large vessels, and only very few tiny vessels were visible around the non-perfusion zones. In comparison with superficial microvessels, the deep microvessels were less damaged with the progression of DR, and microvessel tortuous dilation could be seen around the fovea in patients with mild/moderate NPDR, but the non-perfusion area was not obvious. In severe NPDR, some non-perfusion areas could be seen, but the area was small and the distribution was relatively scattered. The non-perfusion areas in PDR patients were significantly increased, distributed in all quadrants, but mostly concentrated around the fovea, and were divided into different zones by the surrounding microvessels.

## 4. Discussion

Using OCTA scans, patients’ retinal vessels could be stratified into superficial capillary plexus (SCP) and deep capillary plexus (DCP), where the SCP is positioned to extend from the inner boundary membrane to the upper 10 μm of the inner plexiform layer, and the DCP is positioned to extend from the 10 μm above the inner plexiform layer to the 10 μm below the outer plexiform layer. In this study, we compared the vessel density in different regions at the SVD and DVD levels in patients with different stages of DR. The differences were statistically significant, and the VD of each region of the SVD, except the central region, was lower in patients with mild/moderate DR than in patients with NDR. The VD of patients with severe/PDR was lower than that of patients with mild/moderate NPDR; and the VD of each region of the SVD, except the fovea was moderately negatively correlated with the progression of DR disease. This suggested that as DR disease progressed, its superficial vessel density gradually decreased. As DR progressed, the VD in the macula continued to decrease^[9]^, and SVD values were a more sensitive indicator^[10]^ of DR detection by OCTA. A decrease in SVD could also be observed prior to the diagnosis of DR by imaging^[11]^. Zhang et al^[12]^. suggested that microvascular damage in the macular periphery of the retina could be observed in patients with NDR, before the appearance of DR signs such as microaneurysms, which might be one of the reasons for the continued decrease in VD in DR patients.

In previous studies, it was controversial whether deep vessel was lower in patients with PDR and NPDR than in patients with NDR, and different studies have differed with regards to the relationship between deep vessel density and DR progression^[13]^. In the present study, the VD of each region of DVD was slightly higher in patients with mild/moderate NPDR than in patients with NDR; except for the in the fovea, the VD of each region of DVD was lower in patients with severe/PDR than in patients with mild/moderate NPDR, and the difference was statistically significant, indicating that the VD of each region was lower in patients with severe/PDR than in patients with mild/severe NPDR, but the correlation analysis with DR progression was all low. Superficial vessel or involuntary rotation of the patient’s eye during the calculation of deep retinal vessel density by the instrument might affect the detection of deep vessel and produce measurement errors^[14]^. Interstitial refraction might also be a contributing factor to the different results of deep vessel density in different studies^[15]^. In the present study, most of the patients included were elderly, and refractive interstitial clouding, caused by cataract and vitreous opacification may be one of the factors affecting the results of DVD measurements and their correlation results with DR progression^[16]^.

OCTA is a safe and easily reproducible examination method widely used to observe retinal microvascular changes in DR patients. It is a reliable tool for clinical diagnosis and assessment of the condition of DR patients and is a major hotspot for clinical research. Samar et al.^[17]^ concluded that the vessel density of DR patients was positively correlated with their LogMar visual acuity, and Tsai et al.^[18]^ found by correlation analysis that the VD of DR patients was positively correlated with the corresponding regional retinal sensitivity. Previous studies have shown that VD in the macula tended to decrease, as the course of DR progressed. However, their studies mostly tended to compare the variability between VD outcomes in patients with different conditions rather than the correlation between VD and DR progression. There was also a lack of large population studies of different ethnicities collected for the OCTA device to provide standardized data on individual metric measurements as a reference range to use VD alone to make judgments about patient DR staging in clinical work^[19]^. In the present study, although the difference in VD was statistically significant at the DCP level in patients with severe or PDR in comparison with other groups, the correlation was only a low correlation. In contrast, there was a moderate correlation between SCP and DR progression. This showed that a simple difference analysis was not sufficient to show the importance of a single indicator in the analysis of DR progression, and that it was necessary to keep a dynamic eye on the changes in VD of patients. It became particularly important to observe the correlation between VD and DR progression. On the other hand, due to differences in software algorithms, different manufacturers’ instruments yielded significantly different results. The same instrument using different versions of the calculation software might also produce different results. These were the factors that limited the ability of OCTA to observe the progression of DR. More in-depth studies on the role of OCTA in observing the microvasculature of DR are needed.

The foveal avascular zone (FAZ) in healthy eyes is an area delineated by a combination of superficial capillaries, deep capillaries, and is typically round or oval in healthy individuals. The measured area is generally proportional to age^[20]^, usually between 0.231 and 0.280 mm^[21]^. Quantitative parameters such as FAZ area, roundness, and stratified vessel density could be obtained by OCTA scanning. Diabetes mellitus can also cause increased FAZ area, greater circumference, and decreased roundness^[22,23]^, as well as decreased roundness of the peri-macular vascular arch^[24]^. Marques et al. suggested that progressive capillary damage in the macula may be responsible for the larger FAZ^[25]^. Unlike the results of previous studies, the present study analyzed the correlation between FAZ, PERIM and DR progression. Differences in FAZ were not statistically significant with the progression of disease, and PERIM did not correlate with DR progression. The correlation coefficients of AI, FD and DR progression were 0.280, −0.494 (*P*<0.05), respectively, indicating that as DR progressed, the central non-perfusion zone of patients had a more irregular oval shape and the peripheral vessel density was lower. In practice, to avoid the bias caused by manual measurement, we used a computer to automatically calculate the FAZ area. The presence of hard exudate or large tortuous capillaries around the FAZ in patients with severe, PDR led to a small calculation of FAZ area by the software, which might be an influencing factor in the FAZ and PERIM results independent of DR progression.

In combination with the OCTA angiogram (Figures 1 and 2), it could be seen that as DR progressed, patients gradually lost capillaries in the macula and the area of the non-perfusion zone gradually became larger. In patients with severe NPDR and PDR, large areas of non-perfusion were seen. In clinical practice, the VD of the patient’s macular area, especially the SVD, can be used as an indicator to evaluate the progress of DR; and when the VD of DR patients continued to decrease, it might indicate the progression of the patient’s disease. In conclusion, the results of this study showed that OCTA could display the vascular structure and vessel signaling of the retina, clearly demonstrate the changes in microvasculature, detect the subtle changes of retinal vessels in diabetic patients earlier, detect the changes of VD, and FAZ morphology, and could assist in predicting disease progression, staging and prognosis. In clinical applications, the retinal vascular condition of patients could be examined more frequently, and more stringent monitoring of retinal vascular status could be achieved^[26]^.

**Figure 1:**
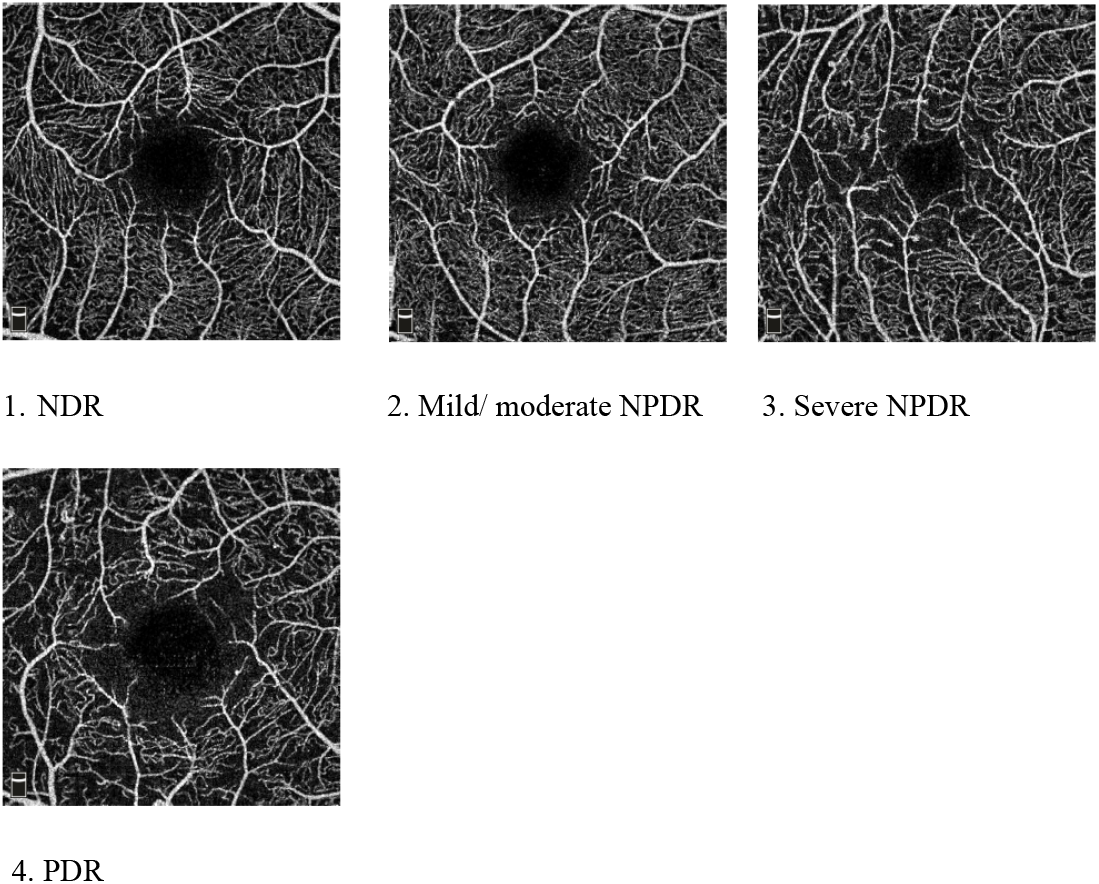
Superficial retinal blood flow imaging in patients of various DR stages.

**Figure 2:**
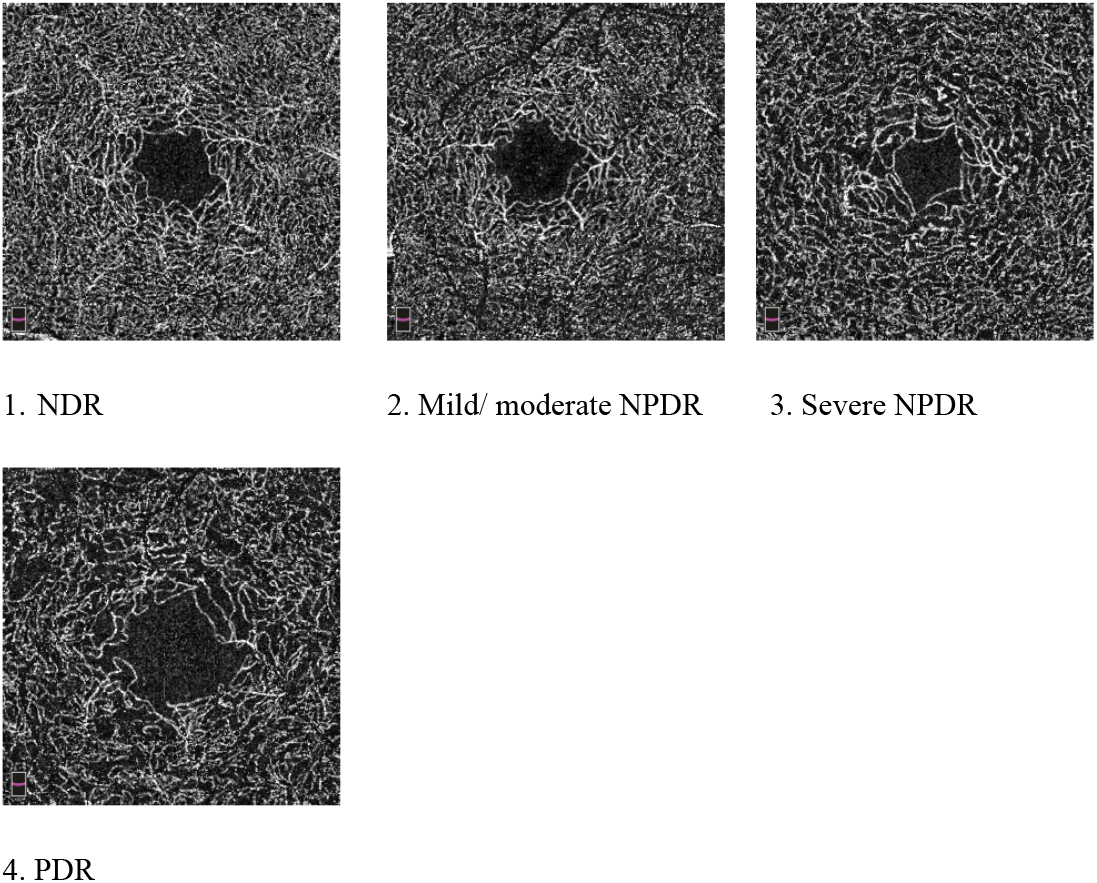
Deep retinal blood flow imaging in patients of various DR stages.

